# Trophic transfer and bioaccumulation of nanoplastics in *Coryphaena hippurus* (Mahi-mahi) and effect of depuration

**DOI:** 10.1101/2024.08.05.606698

**Authors:** Preyojon Dey, Terence M. Bradley, Alicia Boymelgreen

## Abstract

Ocean plastic pollution is a global concern, exacerbated by the distinctive physiochemical characteristics of nanoplastics (NPs), making it crucial to study the impacts on marine animals. While most studies focus on the impacts of waterborne NP exposure, trophic transfer is another key transport mechanism that may also provide insight into the potential transfer of NPs to humans through the food chain. This study investigates polystyrene NP transfer to *Coryphaena hippurus* (mahi-mahi) larvae, a widely consumed fish and significant marine predator, during the early life stage. Using a two-step food chain, *Brachionus plicatilis* (rotifers) were exposed to NPs, and subsequently fed to *C. hippurus* larvae, with exposure durations ranging from 24 to 96 h. Significant NP transfer was observed via the food chain, varying with exposure duration. A depuration study over 72 h, simulating environmental intermittent NP exposure, revealed substantial NP excretion but also notable retention in the larvae. Biodistribution analysis indicated that most NPs accumulated in the gut, with a significant portion remaining post-depuration and some translocating to other body parts. Despite no significant effects on body length and eye diameter during this short study period, histopathological analysis revealed intestinal tissue damage in the larvae.

**Graphical abstract:** 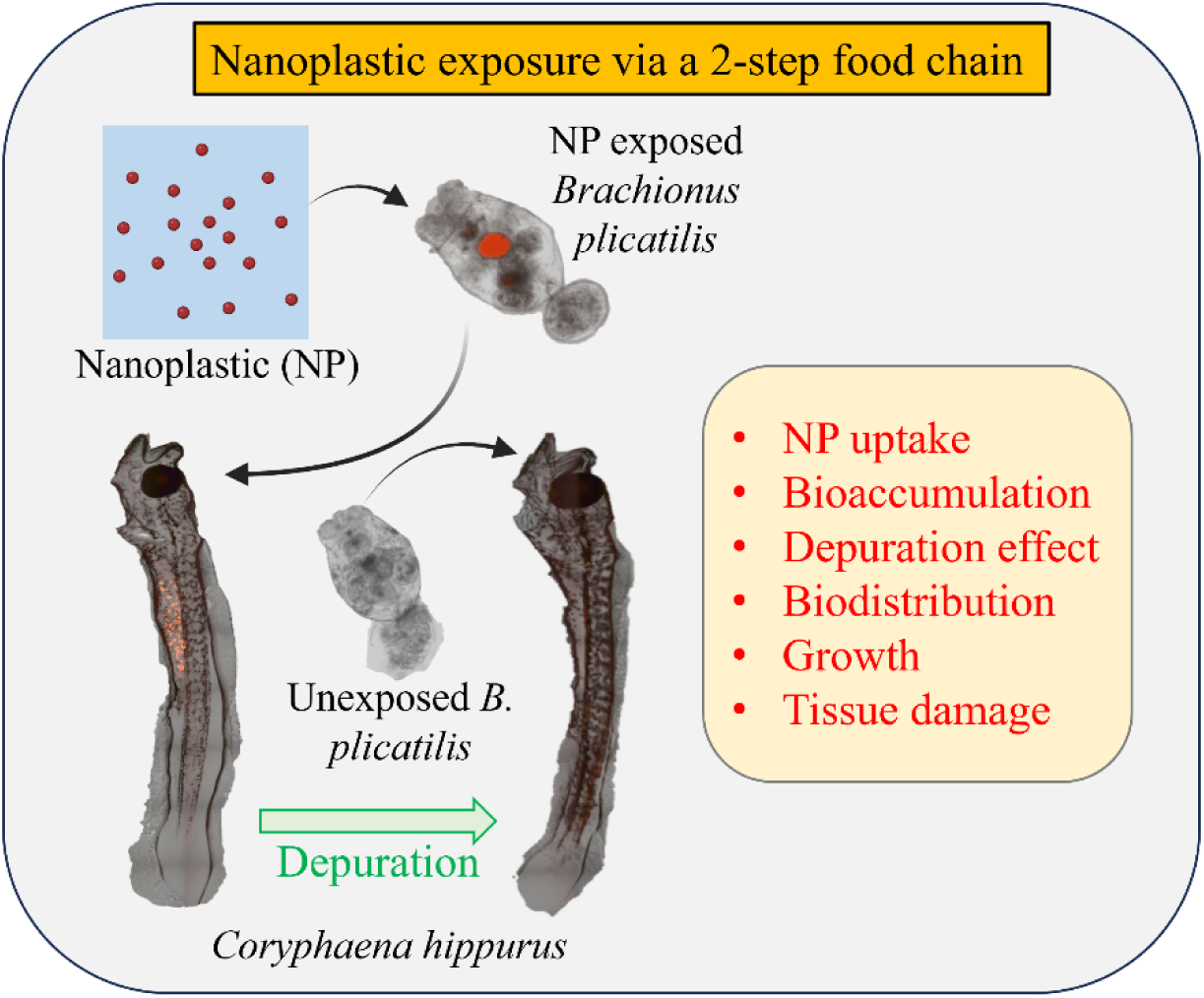

## 1. Introduction

Plastics are extensively utilized throughout a wide range of applications owing to their cost-effectiveness, lightweight nature, and enhanced properties. However, a mere 9% of the plastics generated undergo recycling, leading to a significant proportion of plastic waste being disposed of in our surroundings ^1^. Consequently, a fraction of 3% of the total produced plastic is deposited into the ocean by various pathways such as rivers, runoffs, and direct discharges ^2^, and has emerged as the predominant marine pollutant, comprising 60-80% of the total marine litter ^3^. Prior to reaching the ocean, plastic waste undergoes degradation processes on land, resulting in a range of sizes from macro (5-100 mm) to nanoplastics (NPs) (<1 µm) ^1^. The predominant form of marine plastic waste is macroplastics, accounting for approximately 70-80% of the total ^4^. These macroplastics also undergo degradation and fragmentation in the marine environment through several processes, including photochemical reactions, mechanical forces, hydrolysis, and biological activities, resulting in the formation of secondary microplastics (MPs) and NPs ^4,5^.

NPs pose a significant risk to marine organisms because of their smaller size and weight, as well as their increased surface area ^6,7^, which facilitates widespread dispersion ^8,9^, potential ingestion as food ^10,11^, and higher adsorption and faster leaching ^12,13^. For example, it has been observed that direct exposure to NPs has resulted in adverse effects in marine animals such as embryotoxicity and abnormal gene expression in the sea urchin *Paracentrotus lividus* ^14^; inhibition in hatching ^15^, feeding ^16^, and swimming ^15,16^, molting ^16^ as well as mortality ^15^ in *Artemia*; rapid destabilization of lysosomes, production of reactive oxygen species (ROS) and nitric oxide, and inhibition of phagocytosis in the marine bivalve *Mytilus galloprovincialis* ^16^, etc. NPs have also been observed to be transferred to the upper trophic level via the food chain, such as the food chain consisting of NPs, *D. magna*, and *Carassius carassius* ^17,18^, and to have caused disturbances in feeding, shoaling, and metabolism in upper trophic animals, e.g., fish ^17^. Fish and seafood account for approximately one-fifth of the global animal protein consumption, with an average per capita fish and seafood consumption of approximately 20 kg in 2017 ^19^. It is crucial to understand the retention of NPs in fish, even when they are not continuously exposed to the pollutants, as they can potentially transfer to humans through the food chain. The uptake of plastic or plastic additives and derivatives is associated with various metabolic and functional disorders ^20^, carcinogenicity ^21^, neurotoxicity ^22,23^, obesity ^24^, as well as hemolysis and eryptosis in erythrocytes ^25^ in humans.

This study investigates the ingestion, retention, and distribution of NPs in *Coryphaena hippurus* (mahi-mahi or dolphinfish) larvae exposed to NPs through the food chain for varying durations. It also assesses the effects of NPs on normal growth and potential abnormalities by measuring length and eye diameter, two parameters used in previous studies for similar purposes ^26–28^. Additionally, histopathological analysis is performed to investigate NP trophic transfer-related tissue damage in the intestines. *C. hippurus* is extensively harvested and utilized as a food source for human consumption. Investigation of the potential impacts of NPs on this economically and ecologically significant species is important, particularly during its vulnerable early developmental stage, with a focus on determining the effects of NPs, whether NPs are retained, and the potential implications for human food safety. While there have been studies investigating the effects of environmental disruptions, such as oil spills ^29–32^ and abiotic environmental factors ^33,34^, on *C. hippurus*, there is a lack of research on foodborne exposure to NPs. This route might transfer higher quantities of NPs compared to the waterborne route ^35^, highlighting the need to investigate its implications for this important species. The lower trophic organism employed in this study was the marine zooplankton species *Brachionus plicatilis* (rotifer). *B. plicatilis* has frequently been utilized as a model organism in ecotoxicological investigations owing to its easy availability and cultivation, as well as its rapid growth rate ^36–39^. In this study, polystyrene (PS) NPs were used as a model due to their widespread use in various applications ^40,41^ and their prevalence in marine litter ^42–44^. Polystyrene NPs are also commonly employed in ecotoxicological studies ^15,17,18,45–47^. Following NP exposure, *C. hippurus* larvae were subjected to depuration for varying durations to assess the reversibility of NP retention in the absence of continued NP exposure.

## 2. Experimental

### 2.1 Nanoplastic characterization

PS NPs containing tetramethylrhodamine (TRITC) fluorescent dye with a nominal size of 300 nm dissolved in an aqueous solution at a concentration of 1% solids (w/v) were procured from Thermo Scientific Chemicals (Fluoro-Max, cat. no. R300). Although the size of NPs has been considered less than 100 nm elsewhere ^48,49^, in many studies, plastics less than 1 µm are considered NPs ^50–52^, and the same was considered for this study. Before scanning electron microscopy (SEM) imaging (JEOL FS-100), as-received NP solutions were drop cast onto a double-sided carbon tape affixed to a glass coverslip and air dried overnight. SEM image revealed that PS NPs were spherical in shape (Figure 1A). SEM image analysis shows a size (diameter) distribution of 297.04 ± 18.03 nm (mean ± SD) for the NPs (Figure 1B). Dynamic light scattering (DLS) (Nano ZS, Malvern instruments) results show a Z-average size of 287.24 ± 1.82 nm, a polydispersity index (PDI) of 0.18 ± 0.04, and a Zeta potential of -0.82 ± 0.94 mV, respectively (mean ± SD). PS NPs were suspended in filtered seawater using a vortex mixer (Vortex Genie 2, Scientific Industries) to make NP suspensions with concentrations of 10 mg/L.

**Figure 1.**
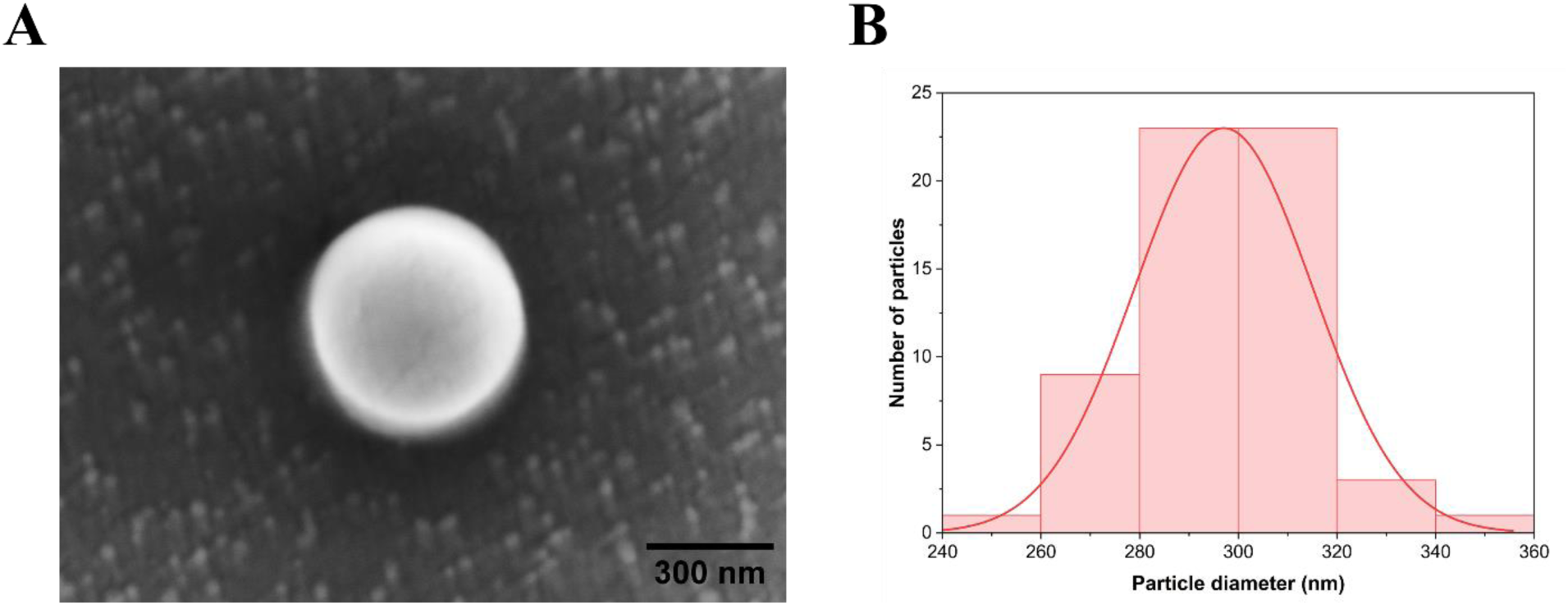
Nanoplastic shape & size. A) Scanning electron microscopy (SEM) images reveal the shape of the polystyrene nanoplastics used in this study (scale bar = 300 nm). B) Size (diameter) distribution of the nanoplastics from SEM image.

### 2.2 Nanoplastic exposure

Approximately 150,000 *B. plicatilis* was transferred to a beaker (1 L) containing NP suspension and held for 3 h. These NP-exposed (NPE) *B. plicatilis* were transferred at a concentration of 30 rotifers/mL, to a container with 100 *C. hippurus* larvae. At least five *B. plicatilis* larvae were extracted from the container at each exposure time point (24, 48, 72, and 96 h), euthanized and preserved in a 70% ethanol solution in preparation for imaging. Additionally, samples of the *C. hippurus* larvae exposed to NP-containing rotifers for 24 and 48 h were collected, transferred to a container containing *B. plicatilis*, which did not receive any NP exposure (non-NPE), for depuration purposes for 24, 48, and 72 h prior to being euthanized and collected in a 70% ethanol solution. The water in the depuration container was changed daily. We note that larvae exposed to NPs through dietary intake for 72 and 96 h did not survive thereafter, likely due to prolonged NP exposure, thereby precluding the investigation of depuration effects on these groups. The control group of *C. hippurus* was only exposed to non-NPE rotifers throughout all experiments and collected at the previously mentioned time points.

### 2.3 Fluorescence microscopy, image processing, and histopathological analysis

The larvae samples were carefully placed within an antifade mounting media (Prolong^TM^ Glass Antifade Mountant, Invitrogen), sandwiched between a glass coverslip and slide. Subsequently, these samples were subjected to imaging using a spinning disc confocal fluorescence microscope (Nikon CSU-X1 mounted on Nikon Eclipse Ti2-E) under both fluorescence (TRITC) and bright-field filters. Images were captured at three distinct Z-levels (bottom, middle, and top) to comprehensively scan the entire depth of the larval samples [Figure 2A(i)]. Due to the microscope’s limited field of view at high magnification and the large size of the larvae sample, images of the entire sample were taken in segments and later stitched together automatically using the NIS-Elements program (large image option). Fluorescence and bright-field images from each Z-level were superimposed on each other and merged into a single composite image using ImageJ [Figure 2A(ii)]. Composite images from different Z-levels were stacked and processed into a single image [Figure 2A(iii)], which was subsequently divided into multiple channels (red, green, and blue) [Figure 2B(i)]. The red channel image was further processed by taking only larvae as the region of interest (ROI), converting it into an 8-bit grayscale image, and then applying Yen’s auto-thresholding method ^53^. The fluorescence intensity of this image was quantified using ImageJ. The quantified fluorescence intensity was then normalized against the control group to determine the fold change in fluorescence intensity [Figure 2B(ii)]. Additionally, the length and eye diameter of larvae samples were measured using ImageJ.

**Figure 2.**
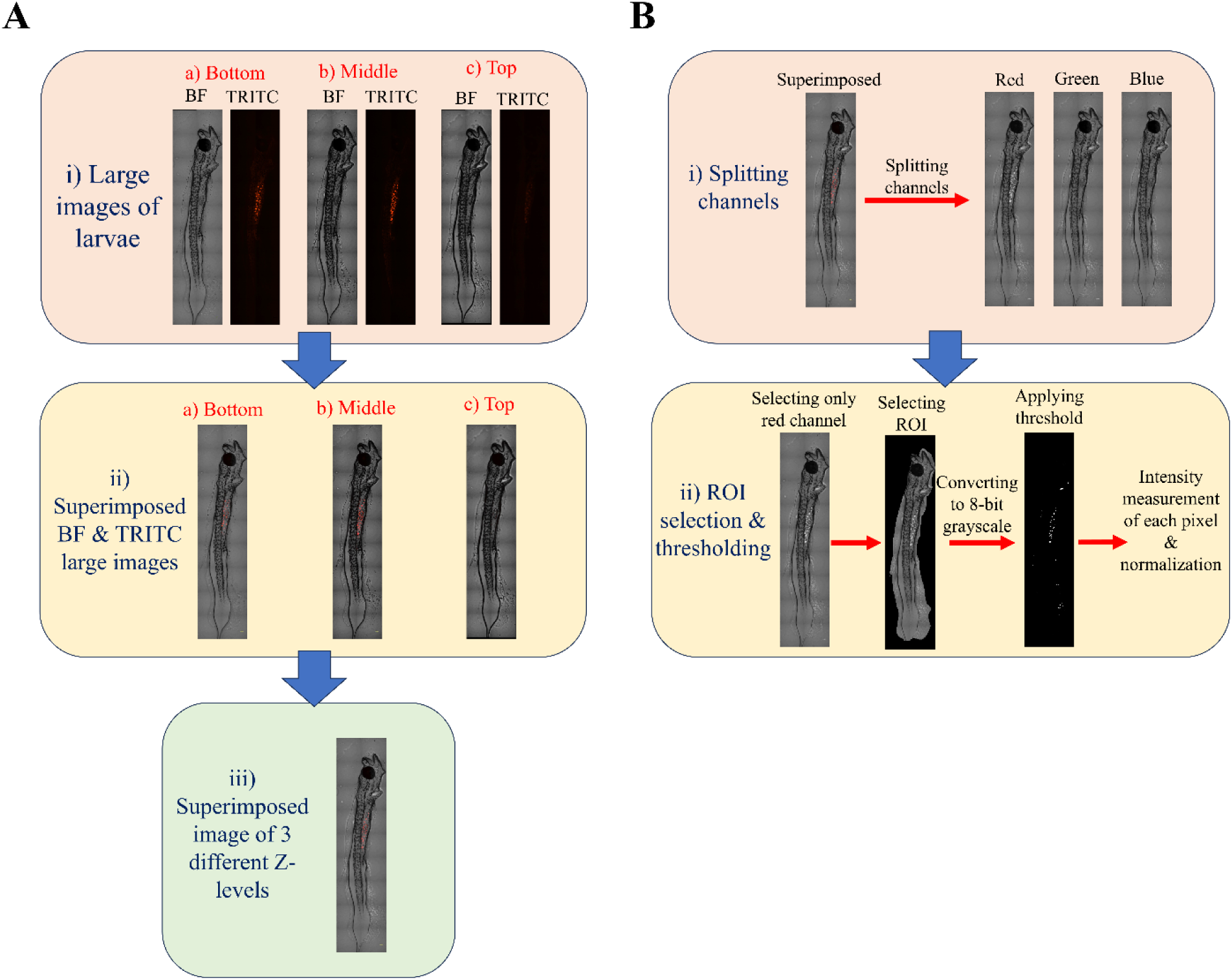
Image Processing and Analysis. (A) Sequential processing of larvae images- i) Acquisition of large images of larvae at various Z levels using different filters (BF: bright-field image and TRITC: fluorescence image), ii) Superimposition of BF and TRITC images of larvae at multiple Z levels, iii) Superimposition of images found from step A(ii). (B) Fluorescence intensity measurement- i) Splitting of the superimposed image from step A (iii) into red, green, and blue channels, ii) Isolation of the red channel image, identification of larvae as the region of interest (ROI), conversion into gray scale image, and application of a threshold and normalization via the control group to quantify fold change in fluorescence intensity.

Histopathological analysis of the larvae intestine was conducted by embedding the samples in paraffin, sectioning at 6 µm, and staining with hematoxylin and eosin (H&E). The glass slide-mounted stained sections were then imaged using bright-field microscopy (Axioscope 5, Zeiss).

### 2.4 Statistical analysis

Statistical analyses were performed with OriginPro (2024b, OriginLab). The data are shown as mean ± standard deviation. Statistical significance was assessed using a one-way analysis of variance (ANOVA), followed by Tukey’s post-hoc test to compare different treatments with the control group. The data was tested for normality using a normal quantile-quantile plot at 95% confidence before ANOVA testing. In all statistical analyses, the results were considered to be statistically significant when the p-value was below 0.05.

## 3. Results and discussion

### 3.1 Effect of different exposure periods on bioaccumulation

Comparison of the superimposed bright field and fluorescence images (Figure 3A) reveals a significant uptake of red PS NPs by *B. plicatilis* within a brief exposure period of 3 h through waterborne exposure. Similarly, images of *C. hippurus* exposed to NPs via trophic transfer (Figure 3B) demonstrate significant uptake and bioaccumulation of NPs, which are absent in the control group. However, as seen by the red NP quantity, the amount of uptake and thus bioaccumulation varied with exposure durations. This observation is corroborated by fluorescence intensity (FI) measurements of *C. hippurus* (Figure 3C), which show a significant increase in FI fold change in the NPE group compared to the control group across all exposure periods, with an average fold change ranging from 18.11 to 581.95 depending on duration. A significant increase in FI with exposure was observed, rising from 24 to 48 h. Although FI decreased at 72 h compared to 48 h, it significantly increased again at 96 h. This pattern suggests a high uptake of NPE *B. plicatilis* and reduced egestion up to 48 h, leading to increased bioaccumulation. Between 48 and 72 h, *C. hippurus* may have egested NPs but ingested fewer NPE *B. plicatilis*, resulting in decreased FI at 72 h. The significant increase in FI at 96 h indicates new uptake of NPE *B. plicatilis* between 72 and 96 h, possibly due to increased egestion of NPs between 48 and 96 h. However, the FI at 96 h exposure is still significantly low as compared to that at 48 h. *C. hippurus* have a high digestive capacity and can feed based on energetic satiation ^54^, and thus should require a large amount of feed as they grow. While the NP-packed *B. plicatilis* were being digested, exposure to PS NPs may disrupt digestive enzyme activities, such as increased levels of lipase, chymotrypsin, and trypsin (possibly due to futile digestion attempts or NP-induced starvation) ^27^ and decreased amylase levels (potentially due to suppressed carbohydrate metabolism) ^27,55^. This could even be exacerbated by the alteration of gut microbiota diversity, potentially leading to dysbiosis, as well as increased levels of cytokines such as *IL-1β*, *IL-8*, *IL-10*, and *TNF-α*, indicating intestinal inflammation, as seen in Zebrafish (*Danio rerio*) exposed to PS NPs ^56^. This could explain the significant decrease in food (*B. plicatilis*) uptake and, consequently, reduced NP bioaccumulation with extended NP exposure duration. Additionally, particle size may influence bioaccumulation trends, as demonstrated in PS NP exposure to Australian bass (*Macquaria novemaculeata*) via trophic transfer ^57^. In this species, uptake and bioaccumulation of smaller NPs (50 nm) increased with exposure duration, while it decreased for larger NPs (1 µm). This trend is likely due to the ability of smaller NPs to cross biological barriers and become internalized in organs more effectively than larger NPs. In this study, the use of ∼300 nm PS NPs, along with their potential increase in size in saltwater suspension due to decreased electrostatic repulsion from high ionic strength ^15^, may also influence bioaccumulation in the larvae.

**Figure 3.**
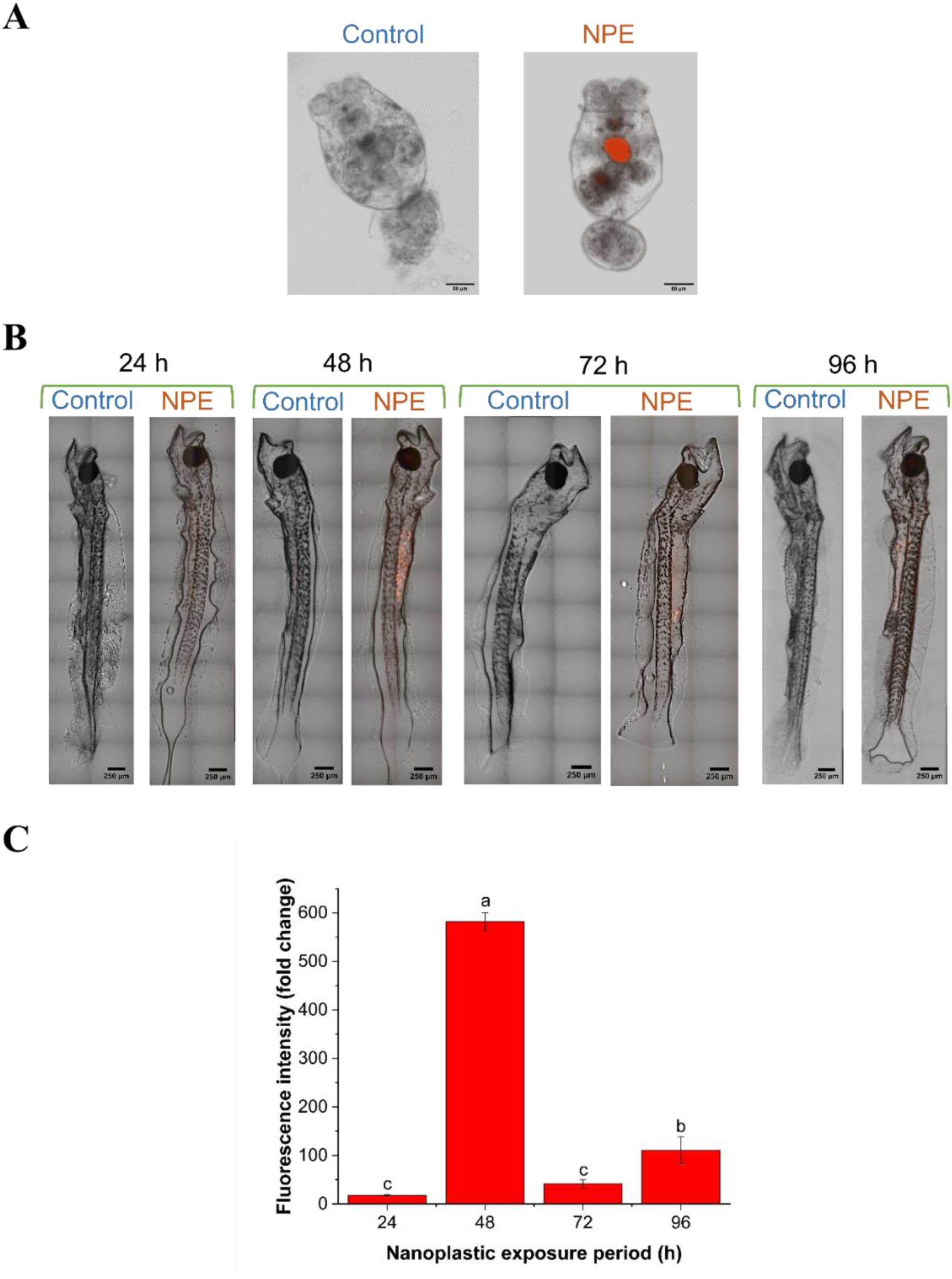
Bioaccumulation of polystyrene nanoplastic in *C. hippurus* through trophic transfer at different exposure periods. Superimposed bright field and fluorescence images demonstrate the presence of red fluorescent nanoplastics compared to the control in (A) *B. plicatilis* following waterborne exposure and (B) *C. hippurus* via trophic transfer after 3 h and various exposure periods, respectively. (C) Fluorescence intensity (fold change as compared to the control) of the red fluorescent nanoplastics accumulated in *C. hippurus* via trophic transfer.

### 3.2 Effect of different depuration periods on bioaccumulation

Figure 4 demonstrates the effect of different depuration periods on the bioaccumulation of NPs in *C. hippurus*, following varying NP exposure durations via trophic transfer. Superimposed bright-field and fluorescence images (Figure 4A) highlight the presence of red fluorescent NPs in *C. hippurus*, absent in the control group, indicating retention of NPs transferred from *B. plicatilis* even after depuration. FI measurements (Figure 4B-left axis) confirm and quantify this retention, showing fluctuating NP levels across depuration periods. Figure 4B-right axis shows the % decrease in FI after a given depuration period compared to the FI at the corresponding exposure period. At 24 and 72 h of depuration after 24 h of exposure, the FI increased unexpectedly as compared to that after 24 h of exposure, resulting in a negative % decrease in FI. This suggests NP ingestion by the larvae during depuration. Despite daily water changes, the breakdown of NPs in feces and subsequent re-exposure via the waterborne route could lead to higher NP accumulation in some larvae. Additionally, possible uptake of NPs by non-NPE *B. plicatilis* from excretion and their consequent dietary intake by C. hippurus may play a role in this process. This raises concerns, as these organisms may act as NP vectors, potentially exposing other organisms via waterborne routes. Other tested conditions showed a significant % decrease in FI, indicating substantial NP excretion and reduced bioaccumulation. For instance, after the highest NP bioaccumulation observed at 48 h of exposure (Figure 3B), the % decrease in FI reached approximately 88% after 24 h of depuration and further reduced to about 98% after 48 h of depuration. However, a significant rise in FI was observed after 72 h of depuration compared to 48 h at the same exposure period, suggesting a similar re-exposure to NPs as previously discussed. NP depuration occurs primarily via feces through a natural excretion pathway ^58^. Despite a significant reduction in ingested NPs after depuration, consistent with previous studies ^55,59,60^, the lowest NP retention (or FI) among all tested conditions (48 h of depuration after 24 h of exposure) was still 8.08 times higher than that of the control in this study. This indicates substantial retention of NPs in *C. hippurus* larvae via trophic transfer even after depuration. Similar substantial NP retention following trophic transfer and subsequent depuration has been reported in small yellow croakers (*Larimichthys polyactis*) ^55^. These results suggest that, in natural environments, where the interaction between NPs and fish is variable, even if fish leave polluted waters, NP retention may persist, ultimately causing negative impacts on them and posing a risk of NP transmission to humans through the food chain.

**Figure 4.**
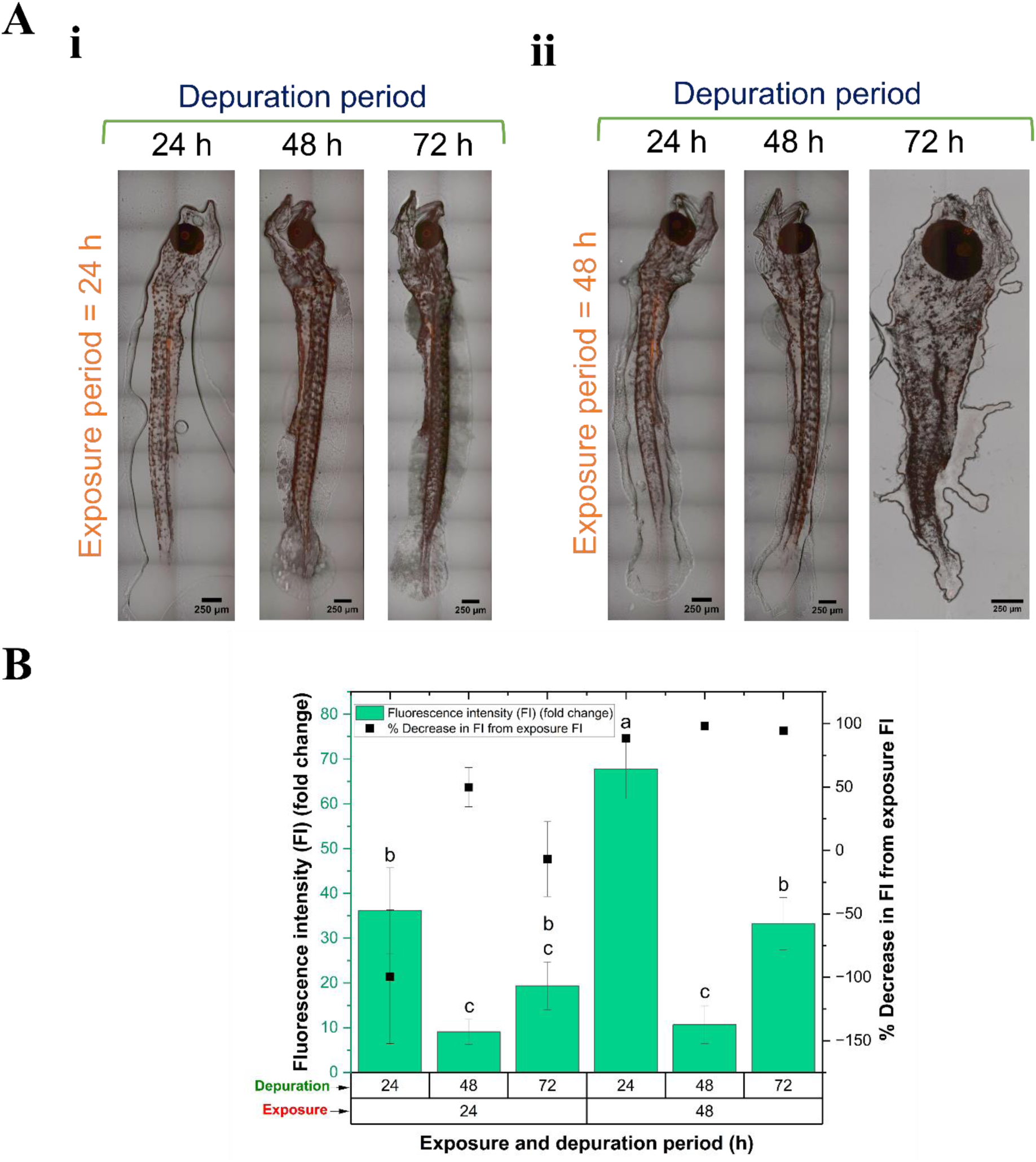
Effect of depuration period on the retention of polystyrene nanoplastics in *C. hippurus* following trophic transfer exposure for various durations. (A) Superimposed bright field and fluorescence images show the presence of red florescence nanoplastics in *C. hippurus* undergoing depuration. (B) Fluorescence intensity (FI) (fold change) of nanoplastics accumulated in *C. hippurus* after various depuration periods compared to the control, and % decrease in FI after depuration from corresponding post-exposure FI.

### 3.3 Biodistribution of nanoplastics

While this study observed the uptake and retention of NPs in *C. hippurus* following various durations of exposure and depuration, it is crucial to investigate whether these NPs transfer to different organs or body parts after entering through dietary exposure. In several previous studies, the biodistribution of NPs was analyzed by dissecting various fish organs and measuring NP accumulation in each ^57,61,62^. However, in this study, we observed NP biodistribution through fluorescence imaging of the whole fish larvae, which did not allow for exact localization within specific body parts. Instead, approximate physiological positions were inferred based on observed fluorescence patterns. Figure 5 shows superimposed bright field and fluorescence images [Figures 5A (i-ii)], and 3D z-stacked fluorescence images [Figures 5B (i-ii), focused on the gut area indicated by the white dotted line in Figures 5A] of *C. hippurus* exposed to NPs via dietary intake for 48 h [Figure 5A(i)], followed by a 72-h depuration period [Figure 5A(ii)]. These conditions were selected as model samples because they represent the highest NP ingestion level and the longest subsequent depuration, as described in sections 3.1 and 3.2, respectively. Since the medium of NP transfer in this study was through food, the highest concentration of NPs was observed in the gut area after NP exposure, as anticipated [Figure 5A(i)]. This observation aligns with previous studies, such as the trophic transfer of PS NPs to Zebrafish via *P. caudatum* ^35^ and *Zacco temminckii* via multiple trophic levels ^63^. This is further illustrated by the 3D fluorescence image of the gut area [Figure 5A(ii)], showing NPs predominantly clustered in distinct lumps. Comparing this with the fluorescence image of NPE *B. plicatilis* (Figure 3A), it appears these NP clusters originate from incompletely digested individual *B. plicatilis* containing NPs in the guts of *C. hippurus*. Upon closer inspection, a significant quantity of NPs had dispersed from these clusters and spread throughout other parts of the gut [Figure 5A(i-d&e)]. Even after 72 h of depuration, during which nearly 95% of NPs were expelled (Figure 4B), a notable amount of NPs persisted in the gut region [Figure 5A(ii-e) & B]. This finding is consistent with the fluorescence intensity results compared to the control group, as discussed in section 3.2. Additionally, a significant amount of NPs was observed around the anus area both after exposure and depuration indicating excretion [Figure 5A-i(f) & ii(b)]. NP accumulation in the gut may be related to mucus secretion, which can trap and immobilize particles ^58^. The retention or excretion of NPs, particularly in the gut, can also depend on their size ^64^, with larger NPs typically taking longer to be excreted ^60^. This effect is expected to be amplified in seawater where NPs tend to aggregate ^65^ and may not readily disaggregate upon ingestion, similar to the disintegration of MPs into NPs, with the behavior potentially depending on the types of plastics involved.^66^. Additionally, NPs can cross biological barriers like the intestinal endothelium due to increased monolayer permeability ^67^ caused by the upregulation of cadherin proteins ^68^, allowing these particles to persist in the body longer and potentially translocate to other tissues via gaps between tissues ^64^. Fluorescence images reveal that some NPs have indeed translocated to various body regions, including areas housing major organs such as the heart, liver, and gall bladder ^69^ [Figure 5A-i(a) & ii(d)], as well as the head [Figure 5A(ii-a)] and caudal peduncle [Figure 5A-i(c) & ii(c)]. NPs were also observed in the fin areas, possibly due to NP excretion from the larvae and subsequent waterborne contact or transit within the fish. Similar translocation of NPs to different organs following foodborne exposure has been observed in previous studies. For example, NPs exposed via the food chain were found to be translocated in the gills, brain, and muscle in *Macquaria novemaculeata* ^57^, and skin, muscle, gills, and liver in *Aphaniops hormuzensis* ^70^.

**Figure 5.**
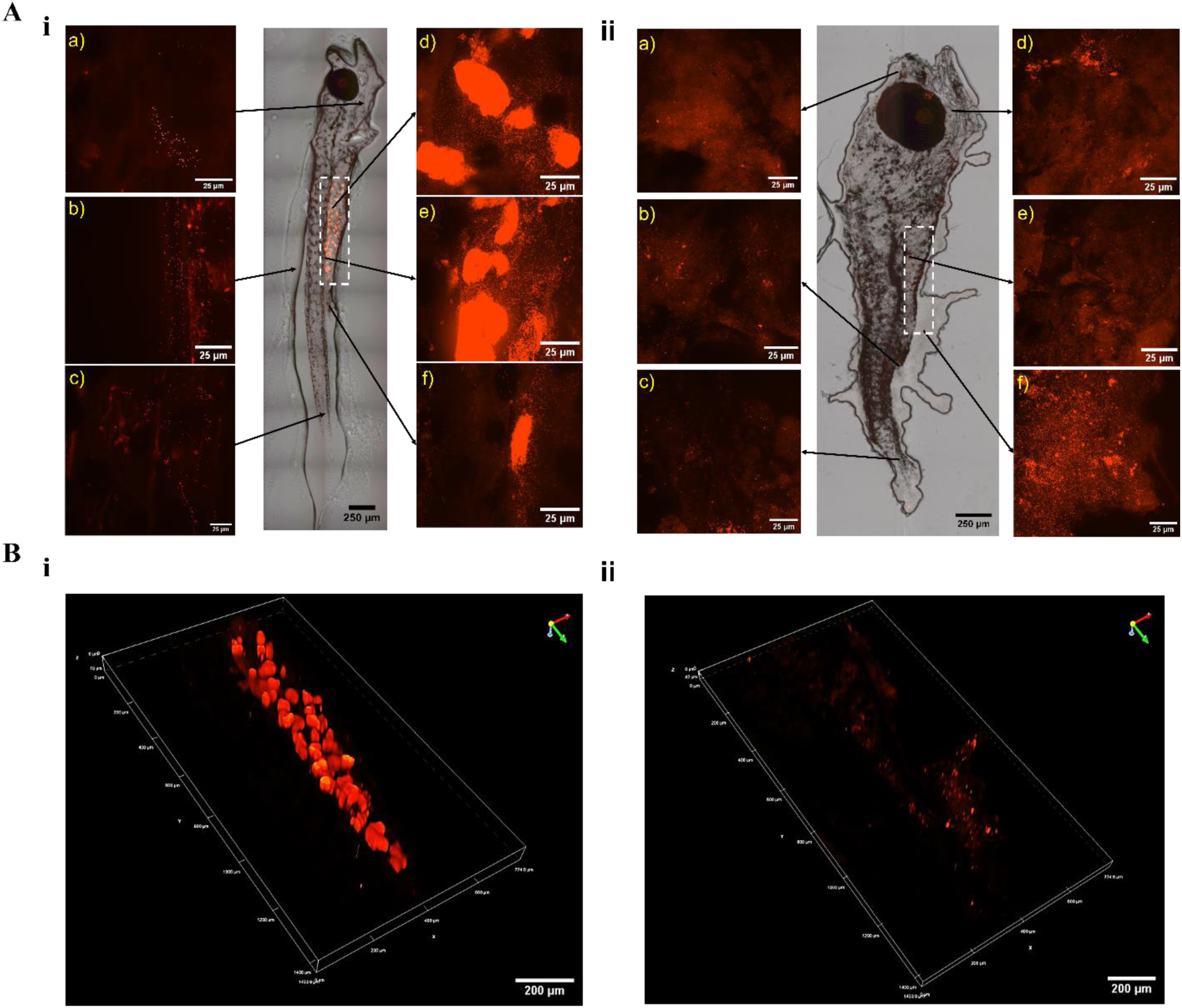
Fluorescent and bright-field images showing the biodistribution of nanoplastics in *C. hippurus* after trophic transfer and depuration. A) Superimposed fluorescent and bright-field images of *C. hippurus* after (i) 48 hours of NP exposure via dietary intake, followed by (ii) a 72-hour depuration period. B) 3D z-stacked fluorescent images of the gut area indicated by the white dotted line in Figure 5A (i & ii).

### 3.4 Effect of nanoplastics on growth

Fish length and eye diameter have been investigated in previous studies to assess the impacts of NPs on growth and potential abnormalities ^27,71^. Figure 6 illustrates the effect of NPs and depuration on the length and eye diameter of *C. hippurus* larvae in our study. To account for age-related variations, larvae of the same age are grouped together for better comparison. The total experimental duration is broken down into control, exposure, and/or depuration phases for clarity. NPs could potentially decrease growth rate (body length) by reducing food intake ^72^. Additionally, exposure to NPs might limit eye diameter by decreasing the number of retinal cells through increased apoptosis, a mechanism also observed with exposure to certain food colorants ^73^. However, in our study exposure to PS NPs through trophic transfer or subsequent depuration did not significantly affect the length of *C. hippurus* larvae under any experimental conditions compared to the control group of the same age (Figure 6A). This lack of effect may be attributed to the relatively short lifespan (maximum 96 h of exposure) investigated in this study. Surprisingly, only the larvae in the 120-h experimental condition (48 h of exposure followed by 72 h of depuration) had significantly shorter lengths than the other experimental groups. However, we note that the same NP exposure (48 h) with lower depuration periods did not show this effect, suggesting it might be unrelated to NP exposure or possibly a delayed impact. Hence, the exact reasons for this observation remain unclear. Similarly, no significant impacts on eye diameter were observed between same-age groups (Figure 6B). As discussed, although some studies have shown impacts of NPs and MPs on the length and eye diameter of aquatic organisms, including fish ^72,74,75^, other studies, found no significant impacts ^27,71,76^. These differences may also depend on factors such as test species, NP type, size, and concentrations ^71^.

**Figure 6.**
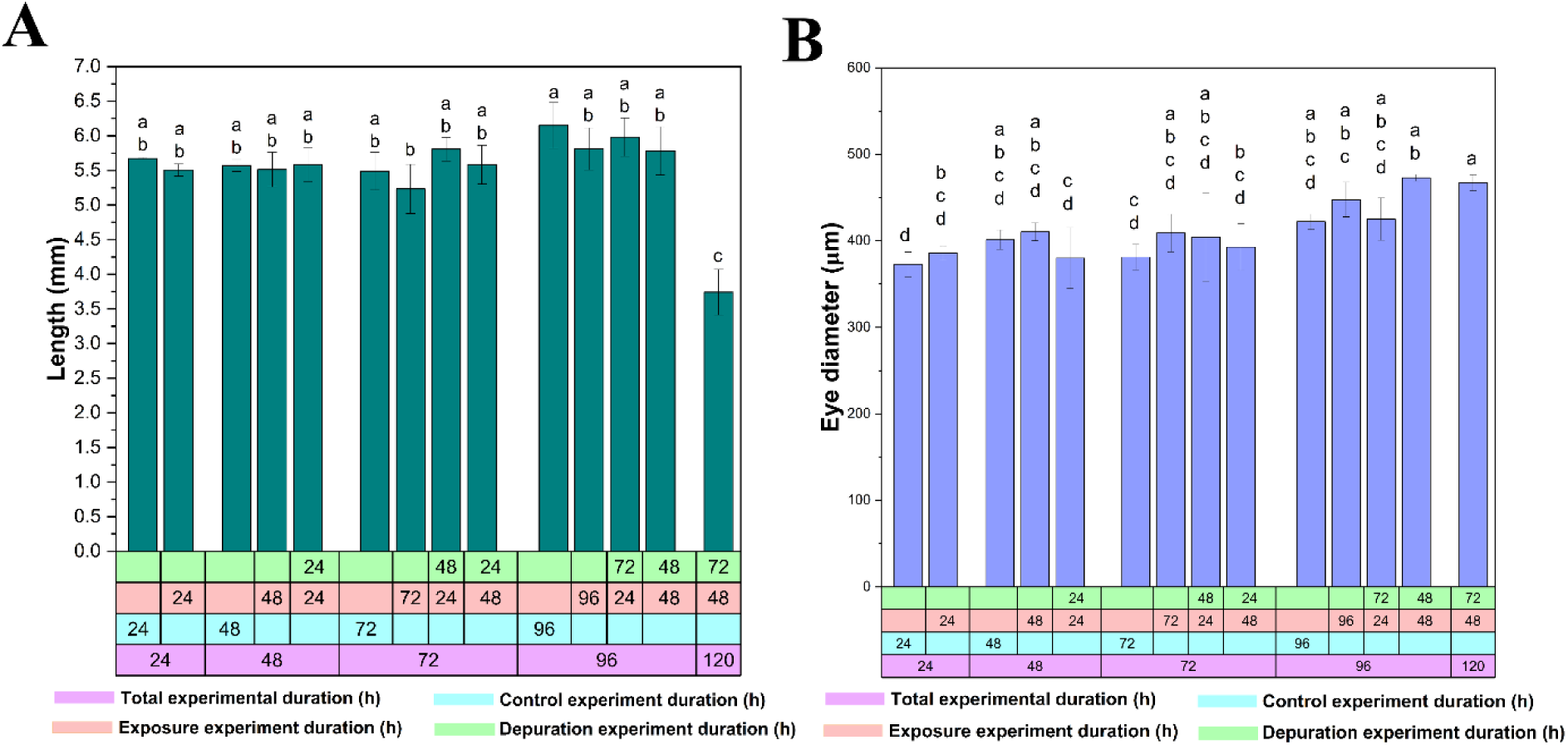
Effects of trophic transfer of nanoplastics on the growth of *C. hippurus* larvae. (A) Length and (B) eye diameter measurements after different exposure durations and subsequent depuration periods.

### 3.5 Histopathological change

As observed in Figure 5 and discussed in section 3.3, most NPs were accumulated in the gut area, warranting investigation of potential tissue damage in the intestine. Figure 7 illustrates the histopathological changes in the H&E-stained intestine of *C. hippurus* larvae following exposure to NPs and subsequent depuration. The images [Figure 7(B-E)] clearly show that NP exposure leads to intestinal injury characterized by shortening, integration, and degradation of villi, regardless of the exposure duration, compared to the control (Figure 7A). The degradation of villi by NPs likely results in villi integration to reduce the contact area between the pollutants and tissue ^77^. Even after 72 h of depuration following 24 h of exposure, similar impacts were observed (Figure 7F). This further explains why food uptake dropped significantly with increasing exposure duration, as observed in Figure 3. Comparable intestinal damage has been reported in other fish species exposed to PS NPs, such as largemouth bass (*Micropterus salmoides*) ^78^, zebrafish ^79^, and grass carp (*Ctenopharyngodon idella*) ^80^. NP-induced tissue damage is generally attributed to the generation of ROS and subsequent oxidative stress ^81^, which results from insufficient production of antioxidants like catalase (CAT) and superoxide dismutase (SOD) that act as defense barriers ^82^. However, an excess of these antioxidants can damage essential biomolecules such as DNA and protein ^77,83^.

**Figure 7.**
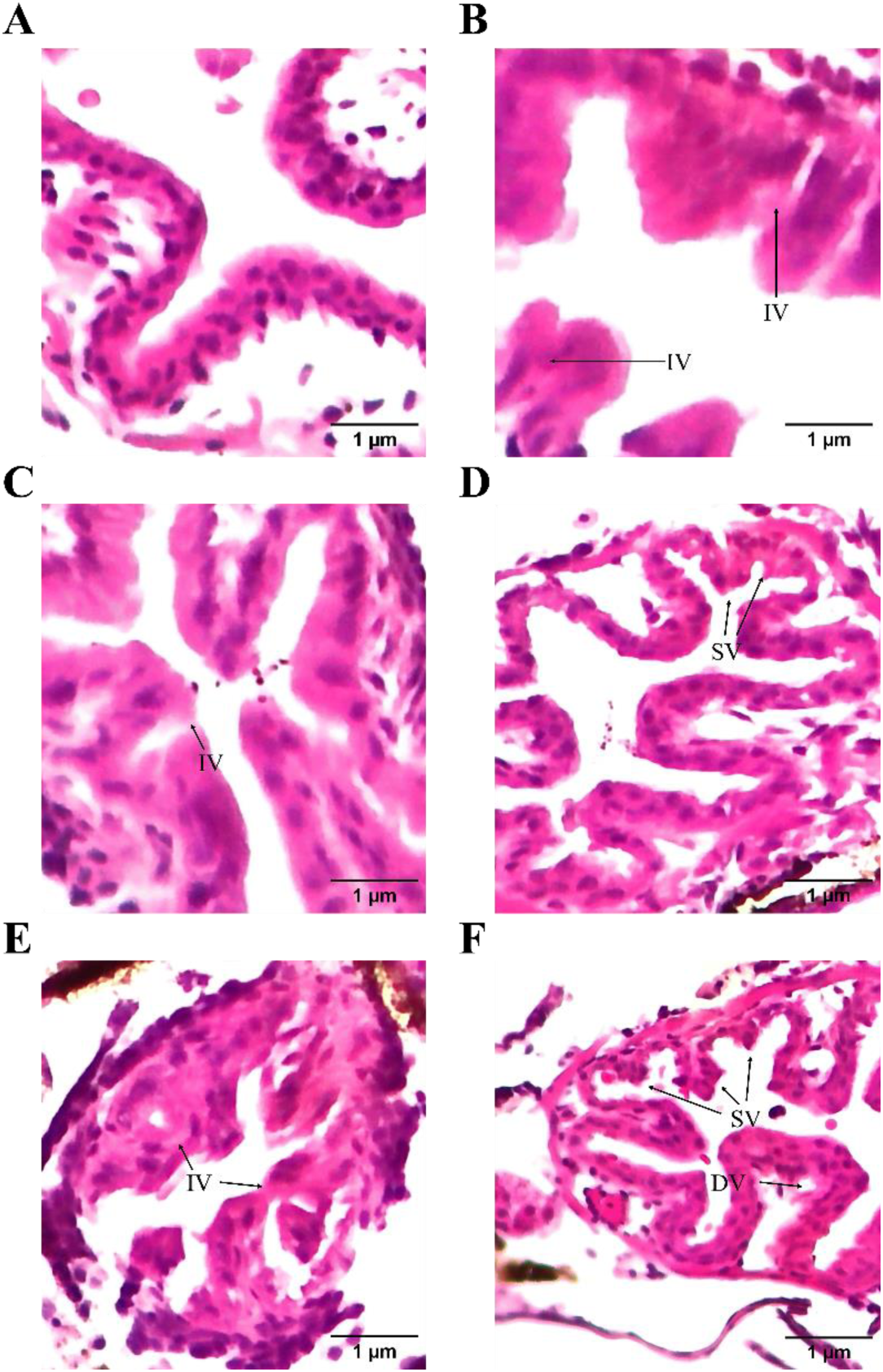
Histopathological changes in the intestine of *C. hippurus larvae* following exposure to nanoplastics via trophic transfer and subsequent depuration. H&E-stained intestine sections of: (A) Control group, and larvae exposed to nanoplastics for (B) 24 h, (C) 48 h, (D) 72 h, (E) 96 h, and (F) 24 h of exposure followed by 72 h of depuration. Integration of villi (IV), shortening of villi (SV), and degradation of villi (DV).

## 4. Conclusion

In this study, we investigated the trophic transfer of polystyrene NPs to the commercially and ecologically significant fish species *Coryphaena hippurus* via a two-step food chain involving *Brachionus plicatilis*. We also examined the effects of depuration on *C. hippurus* when exposed to NP-free *B. plicatilis*. Our findings revealed that NP uptake initially increased significantly with time but later decreased. Depuration significantly reduced ingested NP quantities, yet a substantial portion of NPs remained, indicating implications for fish health and NP trophic transfer to humans. Unexpected increases in NP uptake in some cases during depuration were also observed, potentially due to re-exposure from the feces of other larvae. Biodistribution analysis showed that most NPs accumulated in the gut, forming clusters, while smaller amounts were translocated to other body parts, including the head, caudal peduncle, and areas containing vital organs such as the heart and liver. Even after depuration, significant NP presence persisted in the gut. No significant effect on body length or eye diameter was observed, probably due to the short experimental duration. However, histopathological analysis revealed significant gut damage, including integrated and shortened villi and degradation, regardless of exposure duration or depuration levels. This damage along with potential inflammation ^56^, disrupted digestive enzyme activity ^27^, and altered gut microbiota ^56^ likely contributed to reduced food uptake over time. Over longer durations, these factors could result in significant growth inhibition due to insufficient nutrient absorption.

While this study highlights the negative impacts of NP uptake via the foodborne route in *C. hippurus* larvae, including bioaccumulation, biodistribution, growth effects, and intestinal tissue damage, our analysis was limited to end-point measurements, preventing real-time impact assessment. Future studies could benefit from microfluidic technologies, which allow for fish larvae ^84–86^ and NP exposure ^15^ studies with the possibility of continuous monitoring ^15,87^. Microfluidic integrated sensors ^88,89^, actuators ^90–92^ as well as image processing of moving animals using artificial intelligence ^93,94^, can provide more detailed information on NP impacts and facilitate experimental automation.

## CRediT authorship contribution statement

**Preyojon Dey:** Conceptualization, Methodology, Investigation, Formal analysis, Visualization, Writing - Original Draft, Writing - Review & Editing **Terence M. Bradley:** Conceptualization, Supervision, Formal analysis, Writing - Review & Editing, Funding acquisition **Alicia Boymelgreen:** Conceptualization, Supervision, Formal analysis, Writing - Review & Editing, Funding acquisition.

## Declaration of competing interest

There is no competing interest to declare.

## Acknowledgment

This work is supported by the National Science Foundation (award number: 2038484) and FIU University Graduate School Dissertation Year Fellowship. Graphs were plotted using OriginPro 2024b (https://www.originlab.com/). Schematics were created using Biorender (https://biorender.com/).

## Author Information

## Notes

### Competing Interest Statement

The authors have declared no competing interest.

